# Plastidic Δ6 fatty-acid desaturases with distinctive substrate specificity regulate the pool of c18-pufas in the ancestral picoalga *ostreococcus tauri*

**DOI:** 10.1101/2020.03.10.986216

**Authors:** Charlotte Degraeve-Guilbault C., Rodrigo E. Gomez, Cécile. Lemoigne, Nattiwong Pankansem, Soizic Morin, Karine Tuphile, Jérôme Joubès, Juliette Jouhet, Julien Gronnier, Iwane Suzuki, Frédéric Domergue, Florence Corellou

**Author notes:** Authors contributed equally (Charlotte Degraeve-Guilbault C., Rodrigo E. Gomez, Cécile. Lemoigne). Author Contributions CDG performed the work and analyses on O. tauri (cloning, transgenic screening, HP-TCL, GC-FID, MS/MS); RD performed the work and analyses on N. benthamiana OE (cloning, agro-transformation, FAMES analysis); CL performed most of the lipid analyses of O. tauri and N. benthamiana (agro-transformation, HP-TLC, GC-FID); NP performed the work and analysis on Synechocystis (transformation, screening, TCL, GC-FID); SM designed, performed and analyzed the photosynthesis experiments; KT performed cloning and qPCR experiments; JeJ performed the work on DES localization and qPCR analyses; JuJ performed MS/MS analyses; JG performed the work on DES localization; IS supervised the work on Synechocystis; FD performed and supervised the work on N. benthamiana, helped to organize the MS; FC designed, supervised and performed the research, analyzed the data (O. tauri, N. benthamiana, Synechocystis), wrote the paper. The authors responsible for distribution of materials integral to the findings presented in this article in accordance with the policy described in the Instructions for Authors are: Florence Corellou, and Iwane Suzuki, concerning Synechocystis PCC 6803 lines.

## Abstract

Eukaryotic Δ6-desaturases are microsomal enzymes which balance the synthesis of ω-3 and ω-6 C18-polyunsaturated-fatty-acids (PUFA) accordingly to their specificity. In several microalgae, including *O. tauri*, plastidic C18-PUFA are specifically regulated by environmental cues suggesting an autonomous control of Δ6-desaturation of plastidic PUFA. Sequence retrieval from *O. tauri* desaturases, highlighted two putative Δ6/Δ8-desaturases sequences clustering, with other microalgal homologs, apart from other characterized Δ-6 desaturases. Their overexpression in heterologous hosts, including *N. benthamiana* and *Synechocystis*, unveiled their Δ6-desaturation activity and plastid localization. *O. tauri* lines overexpressing these Δ6-desaturases no longer adjusted their plastidic C18-PUFA amount under phosphate starvation but didn’t show any obvious physiological alterations. Detailed lipid analyses from the various overexpressing hosts, unravelled that the substrate features involved in the Δ6-desaturase specificity importantly involved the lipid head-group and likely the non-substrate acyl-chain, in addition to the overall preference for the ω-class of the substrate acyl-chain. The most active desaturase displayed a broad range substrate specificity for plastidic lipids and a preference for ω-3 substrates, while the other was selective for ω-6 substrates, phosphatidylglycerol and 16:4-galactolipid species specific to the native host. The distribution of plastidial Δ6-desaturase products in eukaryotic hosts suggested the occurrence of C18-PUFA export from the plastid.

**One sentence summary:** Osteococcus tauri plastidic lipid C18-PUFA remodelling involves two plastid-located cytochrome-b5 fused Δ6-desaturases with distinct preferences for both head-group and acyl-chain.

## INTRODUCTION

Marine microalgae synthesize peculiar polyunsaturated fatty-acids including hexatetraenoic acid, (16:4^Δ4,7,10,13^ HTA), stearidonic acid (SDA, 18:4^Δ6,9,12,15^), octapentaenoic acid (OPA18:5^Δ3,6,9,12,15^) as well as very-long-chain polyunsaturated-fatty-acids (VLC-PUFA) (Khozin-Goldberg *et al*., 2016; Jonasdottir, 2019). Due to overfishing and pollution, microalgae are regarded as a sustainable alternative for the production of health-beneficial PUFA inefficiently produced by vertebrates, such as SDA and docosahexaenoic acid (DHA, 22:6 ^Δ4,6,9,12,15^). However, still little is known about the molecular regulation of PUFA synthesis in microalgae. SDA and γ-linolenic acid (GLA, 18:3^Δ6,9,12^) are the Δ6-desaturation products of α-linolenic acid (ALA, 18^Δ9,12,15^), and of linoleic acid (LA, 18:2^Δ9,12^), respectively. The substrate preference of Δ6-desaturase (DES) is considered as the main switch to direct C18-PUFA flows towards the ω-3 or the ω-6 pathways (Shi *et al*., 2015). The ω-3-desaturation of ω-6 substrates establishes a further link between the ω-6 and ω-3 pathway in lower eukaryotes (Wang *et al*., 2013). FA fluxes between lipids and compartments, including cytosolic lipids droplets, are also known to participate in PUFA-remodeling of structural lipid, though knowledge about these fluxes in microalgae remains scarce (Li-Beisson *et al*., 2015; Li, N *et al*., 2016).

Nutrients and abiotic stresses regulate the PUFA content of cyanobacteria and microalgae (Los *et al*., 2013; Khozin-Goldberg *et al*., 2016; Kugler *et al*., 2019). In particular, C18-PUFA from plastidic lipids are highly remodeled in response to abiotic stresses; in the cyanobacteria *Synechocystis* sp. PCC6803, ALA and SDA synthesis is triggered by chilling while in several species from the Chromista kingdom OPA is either increased or redistributed within molecular species (Tasaka *et al*., 1996; Kotajima *et al*., 2014; Leblond *et al*., 2019). As plants, green microalgae display a high amount of α-linolenic acid (ALA, 18:3^Δ9,12,15^) and further the peculiar FA, 16:4^Δ4,7,10,13^ which is typical of the Chlorophyta phylum (Lang *et al*., 2011). Major microalga classes from the Chromista kingdom, such as haptophytes and dinoflagellates, produce both OPA and DHA which are not synthesized by green microalgae. SDA is usually predominant in those species and required for both the synthesis of OPA found in galactolipids and of ω-3 VLC-PUFA, including DHA (Jonasdottir, 2019; Peltomaa *et al*., 2019).

*Ostreococcus tauri* is an ancestral green picoalga that emerged early after the divergence between Chlorophyta and Streptophyta (land plants) (Courties *et al*., 1994; Chrétiennot-Dinet *et al*., 1995; Leliaert *et al*., 2012). It has the most minimal genomic and cellular organization (Derelle *et al*., 2006). This coccoid cell is smaller than 2 µm (picoeukaryote), lacks cell-wall, flagella, as well as an obvious sexual life (Grimsley *et al*., 2010). However, it displays a large panel of PUFA, including HTA, ALA, SDA, OPA and DHA as main components (Wagner *et al*., 2010). *O. tauri* glycerolipid characterization unveiled a clear-cut allocation of PUFA in membranes, with C18-PUFA prevailing in plastidic lipids, OPA being restricted to galactolipids, and VLC-PUFA exclusively found in the extraplastidic lipids consisting of the betain lipid diacylglyceryl-hydroxymethyl-trimethyl-β-alanine (DGTA) and the phosphosulfolipid phosphatidyldimethylpropanethiol (PDPT) (Degraeve-Guilbault *et al*., 2017). We further showed that nutrient starvation resulted in the increase of ALA at the expense of SDA in plastidic glycerolipids and reverberated in the acyl-CoA pool and in triacylglycerols (TAG) with no significant impact in extraplastidic glycerolipids. The sole Δ6-DES previously characterized from *O. tauri* (and related species from the class *Mamiellophyceae*) uses acyl-CoA instead of acyl-lipid as substrates, in contrast to other Δ6-DES from lower eukaryotes (Domergue *et al*., 2005). We reasoned that the regulation of the plastidic C18-PUFA pool involves uncharacterized acyl-lipid Δ6-DES, rather than a transfer of Δ6-desaturation products from the acyl-coA pool to the chloroplast.

In this work, we identified two novel acyl-lipid Δ6-DES, which were plastid located, accepted plastidic lipids as substrates, and displayed distinctive specificities depending on overlapping substrate features. Together with putative homologs from the Chromista and Kinetosplastida lineages, these Δ6-DES cluster apart from the previously characterized Δ6-DES. The discovery of plastidic Δ6-DES and the impact of their overexpression in *O. tauri* points out the requirement of tight regulation of the C18-PUFA pool in microalgae. (Lee *et al*., 2016; Li, D *et al*., 2016). The strategy used is this study further illustrates how overexpression in several host systems of distinctive glycerolipid composition gives insight into DES substrate specificity.

## RESULTS

### *O. tauri* fatty acid desaturase sequences retrieval and analysis

Thirteen canonical DES sequences were retrieved from genomic and transcriptomic databases (NCBI databases). All sequences were manually checked upstream of the predicted start codon, especially in order to assess the N-terminal (Nt) part of the proteins, and extended ORF were validated by cDNA amplification (Table S1, Table S2). Exception made of the acyl-CoA-Δ6-DES and an unknown DES barely related to sphingolipid Δ3/Δ4-DES, all DES were predicted to contain a chloroplastic target-peptide (cTP) (Tardif *et al*., 2012). Among the seven front-end DES three uncharacterized Δ6/Δ8 fatty-acid-DES retained our attention (Table S1, Fig. 1) (Kotajima *et al*., 2014). One Δ6-DES candidate clustered with acyl-lipid Δ6/Δ8-DES and was closest to the diatom *Thalassiosira pseudonana* Δ8-sphingolipid-DES (Tonon *et al*., 2005). The two other candidates were closely related (49.8% identity, 71.6% similarity) and, together with putative homologs, formed a cluster apart from the typical acyl-CoA Δ6-DES from *Mamiellophyceae* species and acyl-lipid Δ6/Δ8-DES from plants, fungi and worms (Fig. 1A, Fig. S1 to S4). These two candidates had each one homolog in the *Mamiellophyceae* species (exception made of *Bathycoccus prasinos*) (Fig. S2, Fig. S3). More distantly related homologs occurred in microalgae arising from secondary endosymbiosis, i.e., species from the Chromista and Euglenozoa supergroup and not in the green lineage (Fig. S4). All homologous sequences displayed the typical Nt-fused-Cyt-b5 domain found in front-end DES. Three His-Boxes, which are known to be involved in both DES activity and specificity, were conserved (Fig. 1B, Fig. S2 to Fig. S4) (Sayanova *et al*., 1997; López Alonso *et al*., 2003). Consensus motifs emerging from alignments corresponded to xHDYxHGRx, WWSxKHNxHH, and QLEHHFLP with a larger conserved region upstream of this latter motif, and showed clearly divergent amino acid signature compared to the acyl-CoA-Δ6-DES His-Boxes (QHEGGHSSL, WNQMHNKHH, QVIHHLFP) (Fig.1B, Fig. S2 to S4) (López Alonso *et al*., 2003). *O. tauri* acyl-CoA-Δ6-DES was previously characterized in yeast and extensively used for VLC-PUFA reconstruction pathway in various organisms including plants (Domergue *et al*., 2005; Hoffmann *et al*., 2008; Ruiz-López *et al*., 2012; Hamilton *et al*., 2016). However, its activity has never been assessed in the native host; The acyl-CoA - Δ6-DES was therefore chosen as a reference to achieve the functional characterization of the two closely related Δ6/Δ8-DES candidates. According to their genomic accessions, these candidates will be referred to as Ot05 and Ot10 and the acyl-CoA-Δ6-DES to as Ot13 (Table S1).

**Figure 1.**
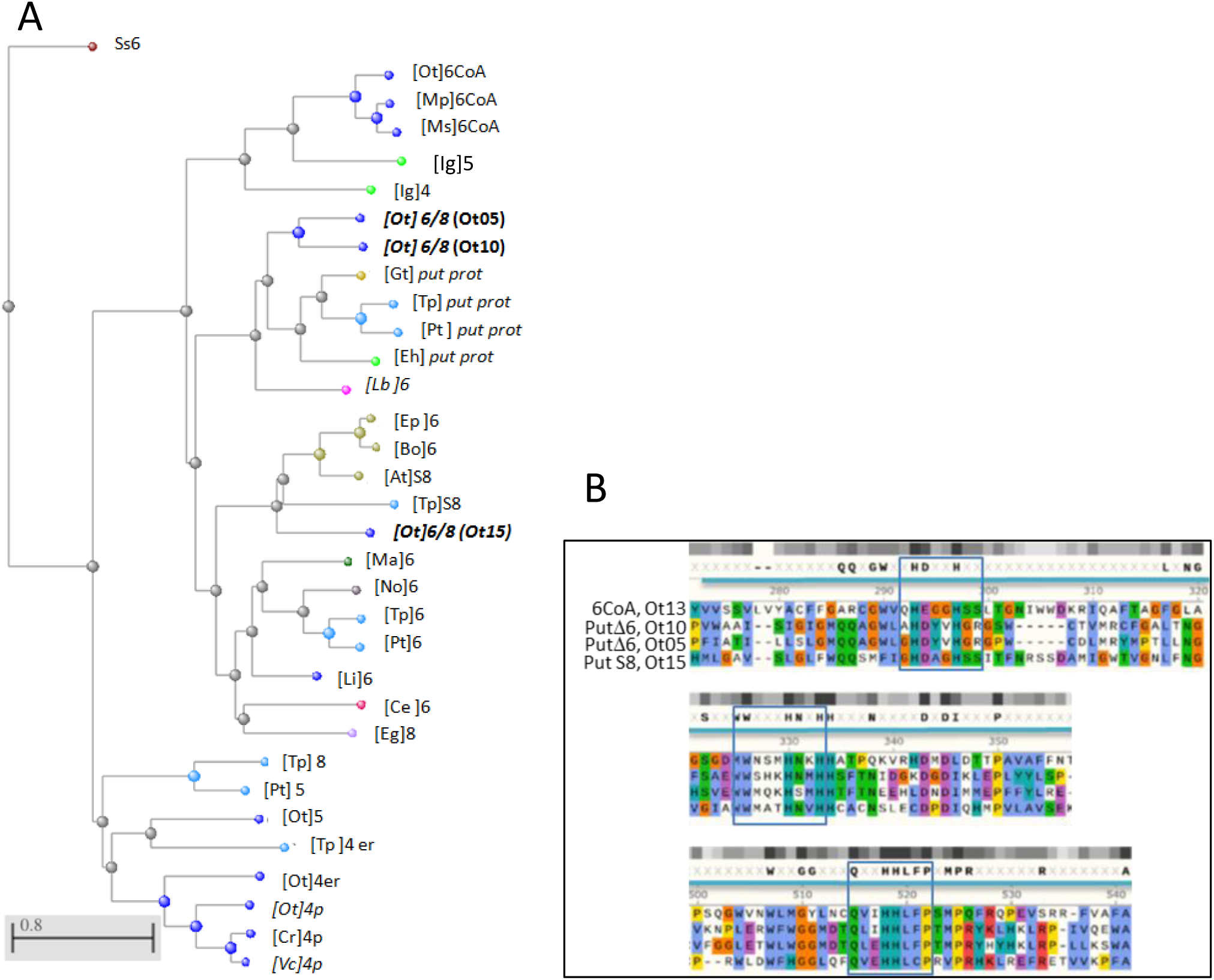
*O. tauri* front-end DES sequence features. **A**. Phylogenetic tree of *O. tauri* front-end DES and closest related homologs (Fast minimum evolution method). Species are indicated in brackets, numbering refers to putative (Italics) or assessed DES Δ-regiospecificty. S, sphingolipid-DES; p, plastidial DES; er, microsomal DES; 6CoA, acyl-CoA Δ6-DES. The three Δ6/8-DES candidates are in bold and their label used in this paper in brackets. Colors of nodes refer to the taxonomic groups: cyanobacteria (purple), eukaryotes (gray), green algae (deep blue), eudicots (beige), cryptomonads (light blue), haptophytes (light green), cryptomonads (yellow), euglenoids (pale pink) kinetoplastids (bright pink), fungi (deep green) nematods (red). [At] *Arabidopsis thaliana*, [Bo] *Borago officinalis*, [Ce] *Caenorabditis elegans*, [Cr] *Chlamydomonas reinhardtii*, [Ep] *Echium plantagineum*, [Eg] *Euglena gracilis*, [Eh] *Emiliana huxleyi*, [Gt] *Guillardia theta* CCMP2712, [Ig] *Isochrisis galbana*, [Lb] *Leishmania brazilensis*, [Li] *Lobosphaera incisa*, [Ma] *Mortierella alpine*, [Mp] *Micromonas pusilla*, [Ms] *Mantoniella squamata*, [No] *Nannochloropsis oculata*, [Ot] *Ostreococcus tauri*, [Pt] *Phaeodactylum tricornutum*, [Vc] *Volvox carteri*, [SS] *Synechocystis* sp PCC6803, [Tp] *Thalassiosira pseudonana*. **B**. Alignment of the acyl-CoA-Δ6-DES and the three Δ6/8-DES candidates in the histidine-box regions. Histidine-box motifs are in blue frames. Color highlighting is based on physical properties and conservation (clustal Omega): positive (red), negative (purple), polar (green), hydrophobic (blue), aromatic (turquoise), glutamine (orange), proline (yellow). Grey blocks highlight conservation only.

### Δ6-DES-candidate localization and activities in heterologous hosts

Full-length ORF or codon-optimized and ORF Nt-truncated, for removing the putative cTP, were expressed in *S. cerevisiae*. Neither the supply of Δ6-substrates nor of Δ8-substrates resulted in the detection of any products. The transformation efficiency was assessed by co-transformation of candidates with the acyl-CoA Δ6-DES, which results in the synthesis of Δ6-products from supplied Δ6-substrates (Fig. S5). We therefore overexpressed full-length proteins in *N. benthamiana* to assess their sub-cellular localization and test their putative Δ6-desaturation activity on the endogenous substrates 18:3n-3 and 18:2n-6. The two putative pΔ6-DES Ot05 and Ot10 fused to the -yellow fluorescent protein (Ct-YFP) exclusively localized at plastids, while the acyl-CoA-Δ6-DES-YFP was at the ER (Fig. 2A). Expression of either fused and non-fused Δ6-DES candidates and acyl-CoA-Δ6-DES resulted in the synthesis of the Δ6-desaturation products 18:3n-6 and 18:4n-3 (Fig. 2B, Fig. S6A). These results unambiguously demonstrated that both Ot05 and Ot10 are plastidic Δ6-DES. A clear trend emerged from the variation of the ω-3 and ω-6 C18-PUFA ratio in individual replicates (Fig. 2B, Fig. S6B).; The 18:4n-3/18:3n-3 ratio was higher in Ot05 overexpressors (Ot05-OE) while the 18:3n-6/18:2n-6 ratio was increased in Ot10-OE and especially Ot13-OE. This suggests that Ot05 preferentially accepted ω-3 substrates while the specificity of Ot10 was higher for ω-6 substrates. The ω-3 and ω-6 C18-PUFA ratio was also used to readily compare the relative impact on glycerolipid classes of each Δ6-DES-OE independently of the overall activity (Fig. 2C, Fig. S7). As a result, extraplastidic phospholipids (phosphatidylcholine PC, Phosphatidic Acid, PA, and phosphatidylethanolamine PE) were mostly impacted in the Ot13-OE (acyl-CoA-Δ6-DES) and the ω-6 ratio 18:3n-6/18:2n-6 was also importantly increased in MGDG. The extraplastidic lipids were affected to a lesser extent in the pΔ6-DES-OE Ot05 and Ot10 (Fig. 2C). Most interestingly, monogalactosyl diacylglycerol (MGDG) was the most altered lipid class in Ot05-OE, while in Ot10-OE, PG showed the highest increase of Δ6-products/Δ6-substrates ratio in Ot10-OE. Impact on sulfoquinovosyl diacylglycerol (SQDG) and digalactosyl diacylglycerol (DGDG) was greater in Ot05-OE compared to the two other OE. Interestingly, disrupting the heme-binding capacity of Cyt-b5 by H>A mutation in the HPGG motif, abolished the activity of both plastidic Δ6-DES (pΔ6-DES) (Sayanova *et al*., 1999) (Fig. S6C).

**Figure 2.**
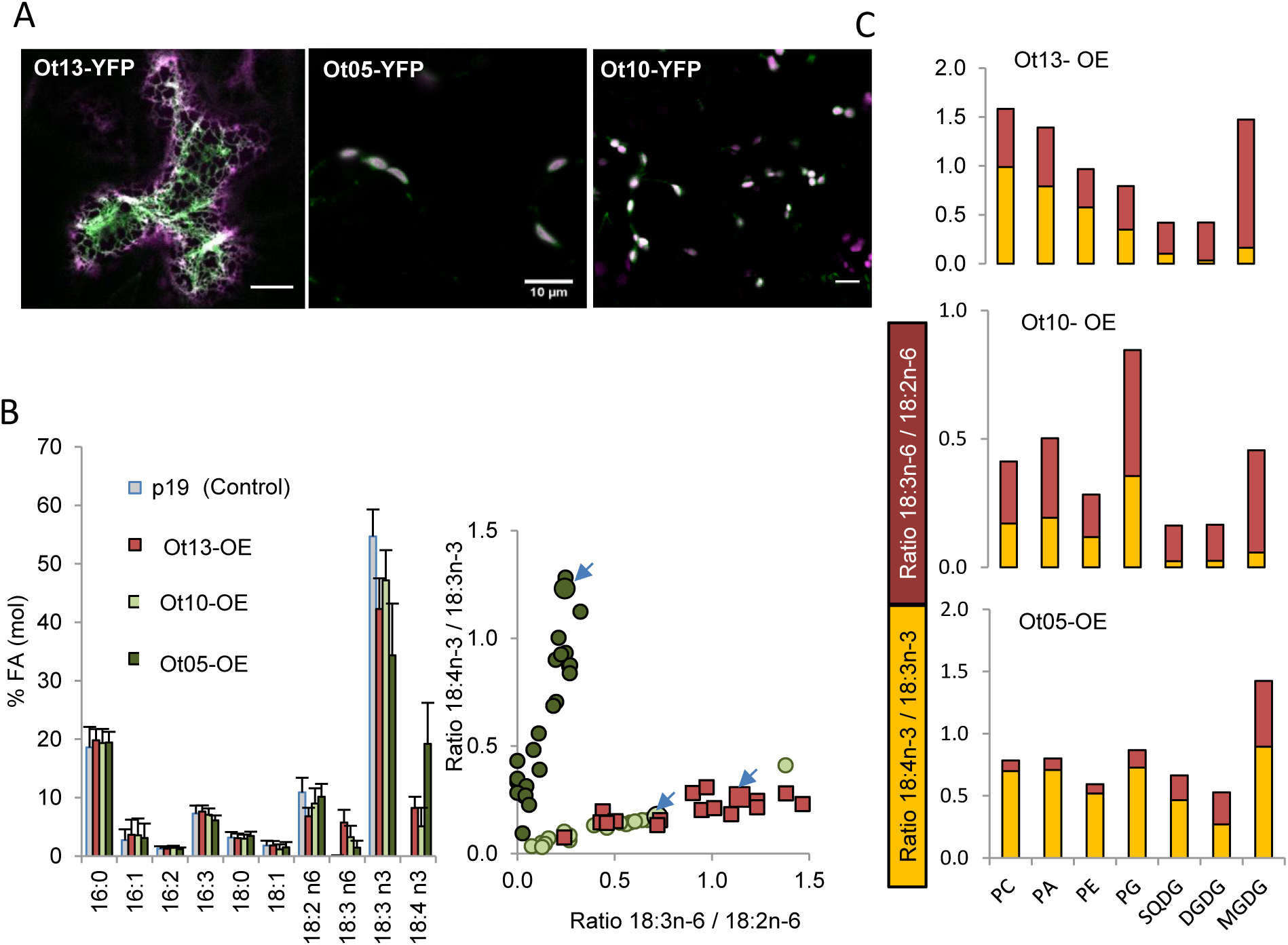
Localization and activities of *O. tauri* acyl-CoA Δ6-DES and Δ6-DES candidates in *N. benthamiana*. **A**. Sub-cellular localisation of transiently overexpressed full-length Ct-YFP-fused proteins. Images merged from YFP chlorophyll or ER-marker (Acyl-CoA-Δ6-DES) fluorescences are shown. Experiments were repeated at least twice. Images represent 100% of the observed cells (n). n=16 for Ot13-YFP (Acyl-CoA-Δ6-DES), n =25 for Ot05-YFP, n=21 for Ot10-YFP. Bar, 10µm. **B**. FA-profiles of DES overexpressors. Means and standard deviations of n independent experiments are plotted as histogram and the relative production of ω-3 C18-PUFA (18:4n-3/18:3n-3) and ω-6 C18-PUFA (18:3n-6/18:2n-6) in each experiment are shown in dot clouds. Dots corresponding to leaves used for the lipids analysis showed in C are indicated by blue arrows. Control lines (p19) n=27, Ot13-OE n=17, Ot10-OE n=21, Ot05-OE n=29. **C**. Relative production of ω-3 and ω-6 C18-PUFA in glycerolipids. Cumulative ratio of pmol percent are plotted 18:4n-3/18:3n-3 yellow bars,18:3n-6/18:2n-6 red bars. On representative experiment out of two is shown (Fig. S7).

To further clarify the substrate specificity of pΔ6-DES, the cyanobacterium *Synechocystis* PCC 6803 was used. This organism not only encompasses the eukaryotic classes of plastidic lipids as major glycerolipids but also allows transgene expression from a similar genomic environment (homologous recombination). *Synechocystis* PCC 6803 has a Δ6-DES (*desD*) and a ω-3-DES (*desB*). In wild type (WT), the selectivity of DesD for galactolipids is reflected by the exclusive distribution of 18:3n-6 in galactolipids. The transcription of *desB* is induced at temperatures below 30°C and results in 18:3n-3 accumulation in PG and SQDG, and of 18:4n-3 in galactolipids. Note that all glycerolipid species are *sn-1/sn-2* 18:X/16:0 combinations.

Complementing either Ot05 or Ot10 in *ΔdesD* cells of *Synechocystis* at 34°C (no endogenous ω-3-DES activity) resulted in 18:3n-6 production at the expense of 18:2n-6 (Fig. 3A). Similar to experiment in *N. benthamiana*, overexpression of H/A mutated version of the Δ6-DES resulted in the absence of Δ6 product, indicating that the integrity of the HPGG motif in the Cyt-b5 domain is required. Ot05 overexpression restored the WT FA-profile, while Ot10 overexpression relatively weak effect. Most interestingly, accumulation of 18:3n-6 occurred not only in galactolipids but was also detected in SQDG for Ot05-OE and in PG for both Ot05-OE and Ot10-OE (Fig. 3B, Fig. 3C, Fig. 3D). Noteworthy, 18:3n-6 accumulation in PG for Ot10-OE was twice as high as that for Ot05-OE, and an isomer of 18:2, likely corresponding to the Δ6 desaturation product of 18:1n-9, was further specifically produced (Fig. 3D). Since no acyl-editing unlikely occurs in cyanobacteria, these evidences support that in addition to galactolipids, SQDG and PG are substrates of Ot05, and PG is a preferential substrate of Ot10.

**Figure 3.**
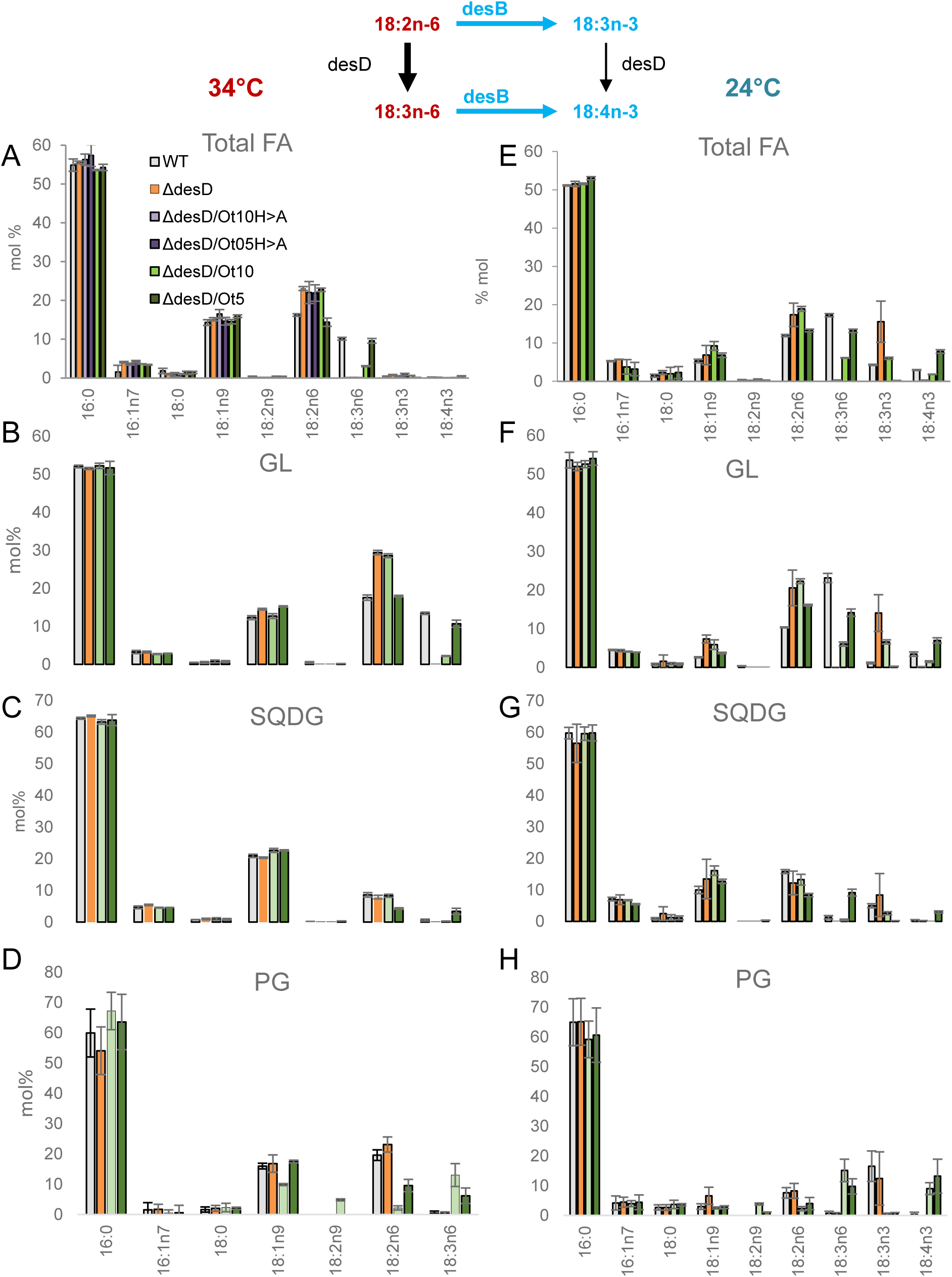
Glycerolipid analysis of Δ*desD Synechocystis PCC6803* Ot5-OE and Ot10-OE. Upper drawing indicates the respective role of desD and desB for the regulation of C18-PUFA in *Synechocystis* PCC6803. C18-PUFA present at 34°C are highlighted in red, those present at 24°C in blue. FA profile of glycerolipids at 34°C (**A, B, C, D**) and 24°C (**E, F, G, H)**. Means and standard deviations of three independent experiments are shown. MGDG and DGDG dsiplayed similar alterations and were cumulated (GL for galactolipids).

*DesB* induction where *Synechocystis* was grown at 24°C led to 18:3n-3 synthesis and further 18:4n-3 accumulation in galactolipids. The increase of 18:3n-6 at the expense of 18:2n-6 reflects that the endogenous Δ6-desaturation of 18:2n-6 is favored and that ω3-desaturation is a limiting step for 18:4n-3 production in WT (Fig. 3E). Overexpression of *O. tauri* pΔ6-DES restored the production of 18:4n-3 in the Δ*desD* background to a similar degree in Ot10-OE as in WT, and to a greater extent in Ot05-OE (Fig. 3E). By comparing the profiles of Ot05-OE with WT, it appeared that 18:3n-6 was decreased to a greater extend and that 18:3n-3 was depleted, indicating that the 18:2n-6, 18:3n-3, 18:4n-3 route prevailed over the 18:2n-6, 18:3n-6, 18:4n-3 route. This is coherent with the results obtained in *N. benthamiana*. Interestingly, 18:4n-3 not only accumulated in galactolipids of both pΔ6-DES OE, but was also detected in PG-species and specifically in SQDG for Ot05-OE (3% of SQDG species) (Fig. 3F, Fig. 3G, Fig. 3H). While 18:X/16:0 galactolipids and SQDG species appeared to be converted more efficiently by Ot05, 18:3n-3-PG seems to be equally well accepted by both pΔ6-DES (Fig. 3D, Fig. 3H).

### Δ6-DES overexpression in *Ostreococcus tauri*

To gain insight into the regulation of C18-PUFA pool by Δ6-DES in the native host, *O. tauri* overexpressors of each Δ6-DES were created using the pOtOXLuc vector (Moulager *et al*., 2010).

### Screening and selection of Δ 6-DES transgenic lines

Phosphate limitation is required for the maximal activity of the high-affinity-phosphate-transporter promoter (promHAPT) driving transgene overexpression. Furthermore, phosphate limitation has been previously shown to enhance the accumulation of 18:3n-3 at the expense of 18:4n-3 in WT (Degraeve-Guilbault *et al*., 2017). Transgenics for each of the Δ6-DES were screened by luminescence recording of the luciferase reporter gene (promCCA1:Luc) and their FA-profiles were further assessed (Fig. S8). Five transgenics with various phenotypes were selected to ascertain the FA-profile and the transgene expression level (Fig. S9). The transgenics displaying the highest luminescence levels showed the most pronounced alterations regarding the amount of C18-PUFA (Fig. 4). In Ot05 transgenics lines 5-3, 5-4 and 5-5, 18:4n-3 was greatly increased at the expense of 18:3n-3, and a less drastic increase of the down-product 18:5n-3 was observed. These variations on ω-3-C18-PUFA were less pronounced in the best Ot10 transgenics (10-2, 10-5), while the impact of Ot05 and Ot10 overexpression on ω-6-C18-PUFA was comparable (Fig. 4A, 4B and 4C, Fig. S9B). Only minor alterations, consisting mainly of 18:3n-3 reduction, were observed for Ot13-OE (Fig. 4, Fig. S9). Additional changes, such as a decrease of 16:4n-3 and a slight increase of 20:4n-6, were observed in the best pΔ6-DES-OE (Fig. S9, Fig. 4D). One representative overexpressor for each Δ6-DES was chosen (Ot05-5, Ot10-5, and Ot13-5, which displayed a similar level of transgene expression) were further used for detailed analysis (Fig. 4D, Fig. S10A). These overexpressors grew similarly as control lines (Fig S10B).

**Figure 4.**
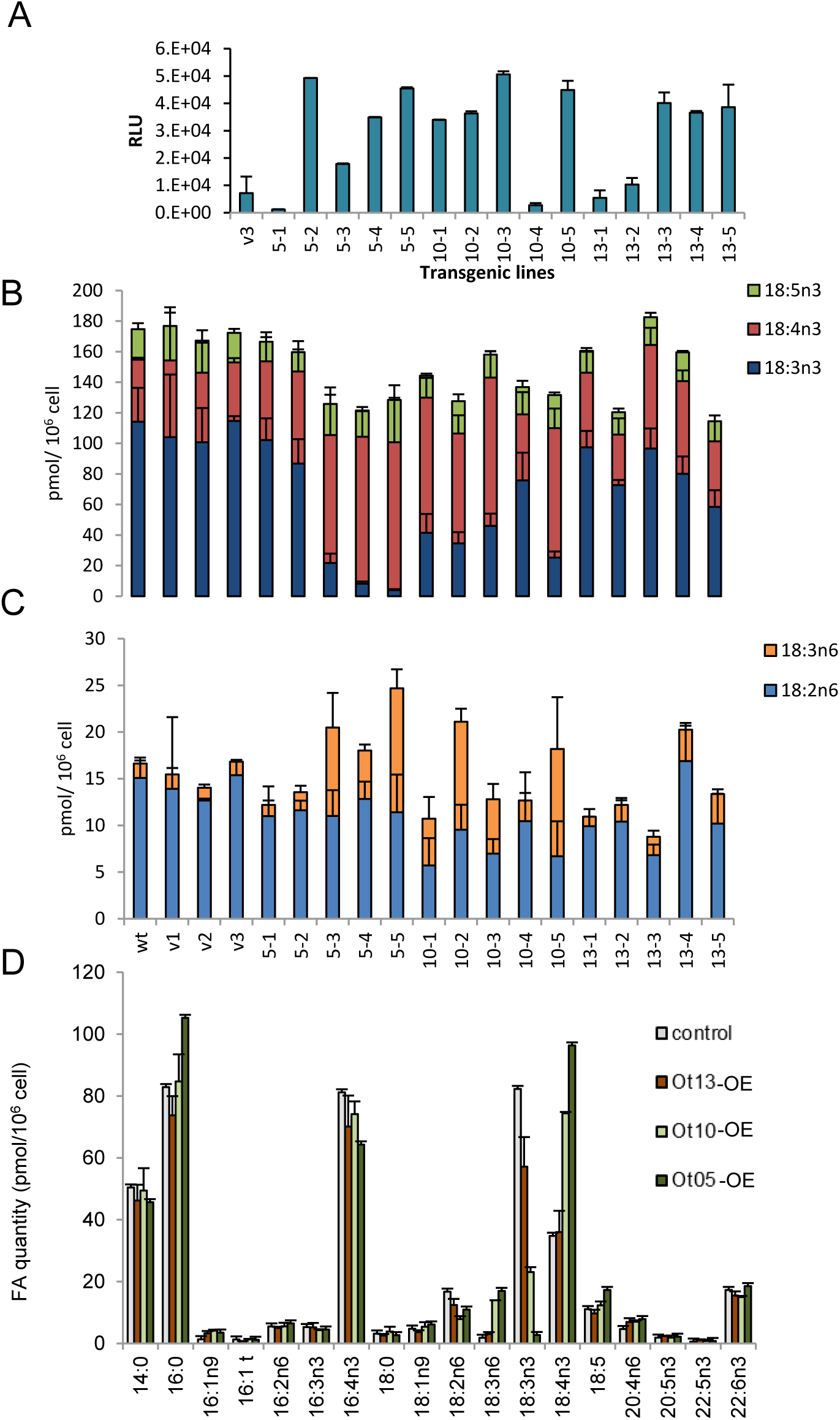
Glycerolipid features of *O. tauri* Δ6-DES-overexpressors. **A**. Luminescence of transgenic lines (Relative Luminescence Units from 200µl). Mean of triplicate and standard deviations are shown. **B** cellular amount of ω-3-C18-PUFA **C**. C18-PUFA cellular amount of ω-6-C18-PUFA. The labels v, 5, 10 and 13 correspond to lines transformed with empty vector, Ot5, Ot10 and Ot13 respectively. **D**. Total glycerolipid FA profiles of lines selected for detailed lipid analysis (0t05-5, Ot10-5, Ot13-5). B to D. Means of triplicate independent experiments and standard deviation are shown.

### Lipidic features of selected Δ6-DES overexpressors

For Δ6-DES OE, minor changes were observed in the proportion of lipid classes consisting mainly of an increased accumulation of TAG, mostly at the expense of MGDG (Fig. S10C). As expected, plastidic lipid FA-profiles were greatly impacted by pΔ6-DES overexpression while only slightly by the acyl-CoA-Δ6-DES overexpression (Fig. 5). Overall FA variations in plastidic lipids followed a similar trend in both pΔ6-DES-OE: ω-6-C18-PUFA were equally impacted in Ot05-OE and Ot10-OE, while the accumulation of 18:4n-3 or of its Δ3-desaturation product 18:5n-3 in 18:5/16:4-MGDG was greater for Ot05-OE (Fig. 5A, B). Ot05-OE also displayed a specific and important decrease of the relative amount of 16:4n-3 in DGDG that was paralleled by an increase of 16:0; however the proportion of 16:4n-3 remained stable in MGDG (Fig. 5A, 5C). Most interestingly molecular species analysis unveiled that the differences between Ot05-OE and Ot10-OE were enhanced for the DGDG species 32:4, 34:4 (18:X/14:0, 18:X-16:0) and the SQDG species 34:4. The relative accumulation of 18:4/saturated fatty-acid combinations in Ot05-OE (ratio to control line) was more than twice as high in DGDG and three times higher in SQDG as for Ot10-OE (Fig. 5D, 5F). In contrast, the variations of C18-PUFA from unsaturated galactolipids species were close to one another in Ot05-OE and Ot-10-OE.

**Figure 5.**
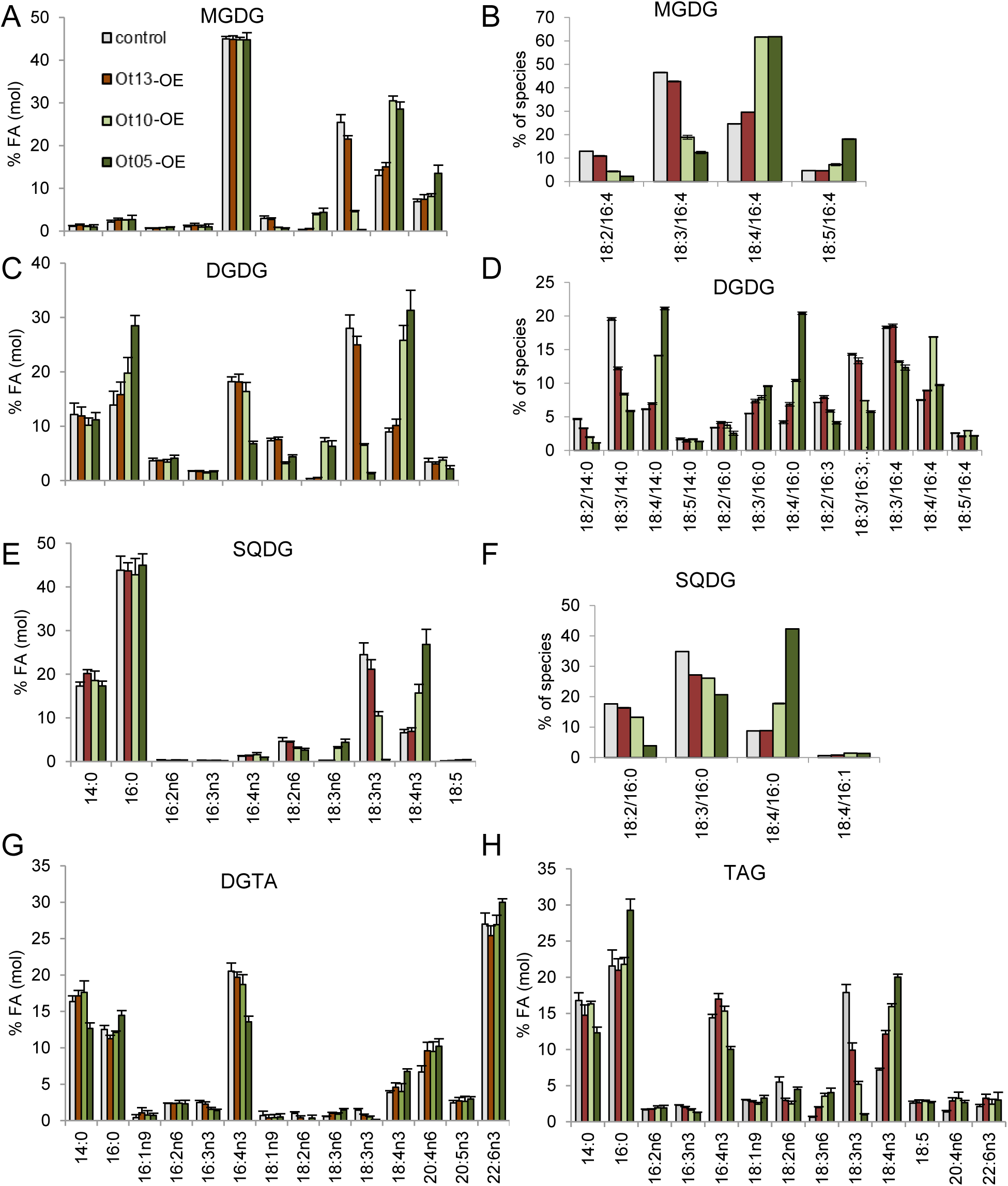
Detailed Lipid analysis of *O. tauri* Δ6-DES overexpressors. **A to F**. Major plastidic glycerolipids. **G, H**. Extraplastidic glycerolipids. For FA-profile analyses, (A,C,E,F,G,H) means and standard deviations of three independent experiments are shown; control line contains the empty vector. **B, D, F**. C18-PUFA molecular species analysis of major plastidic lipids. Means and standard deviations of technical triplicate are shown. Samples used for this analysis are independent from those used for GC-FID analysis; control line is the wild-type (WT).

Extraplastidic structural lipids showed modest alterations consisting mainly of a 30% increase of the proportion of 20:4n-6 in the three Δ6-DES OE (Fig. 5G). Under phosphate limitation, DGTA is the prevailing extraplastidic structural lipid (Degraeve-Guilbault *et al*., 2017) (Fig. S11A, S11B). A specific increase of 18:4n-3 was observed in DGTA for Ot05-OE and was likely relying on the increase of the species 32:4 which includes 14:0/18:4-DGTA (Fig. 5G, Fig. S11C). Similar to the phenotype observed in DGDG, 16:4n-3 was more importantly decreased in Ot05-OE. Several molecular species specifically decreased in Ot05-OE, including 36:8 (putatively 20:4/16:4), 30:4 (14:0/16:4), and 38:10 (22:6/16:4), might account for the overall 16:4 decrease.

Overexpression of each of the three Δ6-DES-OE importantly impacted the FA-profile and molecular species of TAG (Fig. 5H, Fig. S12, S13). Notably for Ot13-OE, the clear alteration of the ratio Δ6-DES-substrates / Δ6-DES-products in the TAG contrasted with the minor alterations detected in structural lipids. As expected, all 18:4-containing species increased at the expense 18:2- and/or 18:3-TAG species in Δ6-DES-OE: for instance, the major molecular TAG species 48:4, 50:4 increased while the major 48:3, 50:3 species decreased (Fig. S12, S13). Noteworthy, the peculiar species 50:10 (16:4/16:3/18:3 and possibly 16:4/16:4/18:2) and 50:11, (16:4/16:4/18:3) were the second most importantly reduced species. As previously described for DGTA and DGDG, the relative amount of 16:4n-3 was specifically decreased in Ot05-OE, while 16:0 was increased (Fig. 5H). Other alterations specific to Ot05-OE consisted of the reduction of the species 48:7 (includes 16:4/14:0/18:3), 50:6 (includes 16:4/16:0/18:2) and 50:7 (includes16:4/16:0/18:3), by about half, and the increase of the proportion of 54:10 (includes 18:4/14:0/22:6), 56:9 (includes 18:4/16:0/22:5) and 56:10 (18:4/16:0/22:6) by more than twice. Therefore, the specific 16:0 increase in TAG from Ot05-OE most likely relies on 56:9 and 56:10; these species are 16:0 *sn-2*-TAG species, i.e., TAG species possibly arising from plastidic DAG precursors (Degraeve-Guilbault *et al*., 2017). Altogether, these observations indicate that FA fluxes toward TAG are differentially affected in the Δ6-DES-OE, and further suggest that 16:4-TAG species, including the peculiar di-16:4 species, are importantly involved in the fine-tuning of C18-PUFA.

### Physiological relevance of pΔ6-DES regulation

Transcriptional regulation of desaturases is known to occur in response to environmental cues. We therefore assessed transcript levels of desaturases involved in the regulation of C18-PUFA pool by phosphate availability, including the putative ω3-DES (Kotajima *et al*., 2014). Consistent with our previous report, the proportion of 18:3n-3 was increased by about one half after the transfer of cells to phosphate depleted medium (Fig. 6A). By that time, the transcript level of Ot05 was decreased by more than 60%, the transcripts levels of Ot10 and of the putative ω-3-DES remained stable while transcripts of the acyl-CoA DES were rather increased. This result indicates that a decrease in Ot05 activity through transcriptional repression results in lowering the 18:4n-3/18:3n-3 ratio under phosphate deprivation.

**Figure 6.**
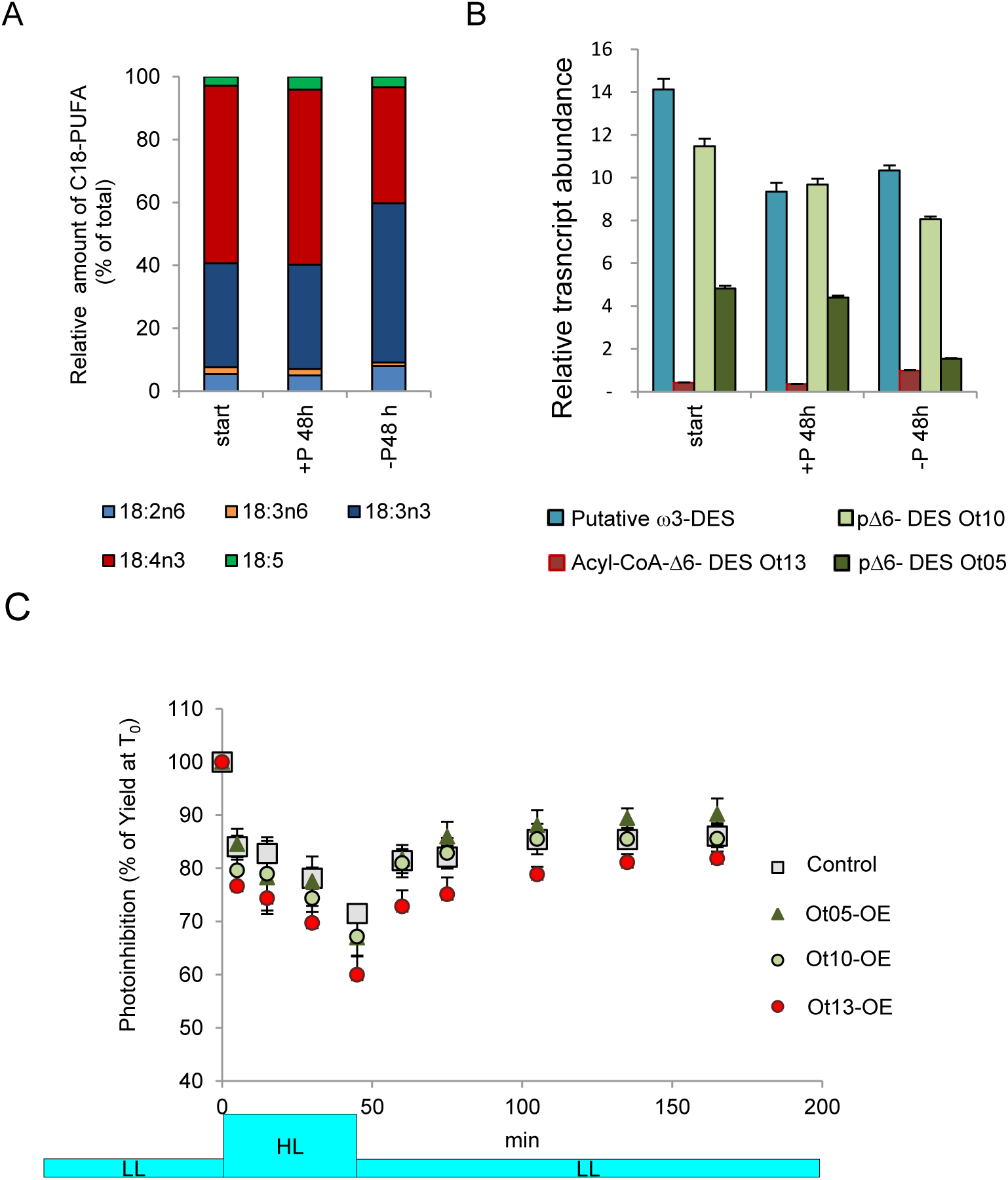
Phosphate limitation and Δ6-DES regulation in *O. tauri*. **A-B**. Impact of phosphate deprivation on C18-PUFA proportion (A) and desaturases transcript levels (B). **C**. Photosynthetic inhibition responses of *O. tauri* Δ6-DES-OE in phosphate-limited conditions. Photosynthesis efficiency (Yield) was assessed under 30 µmol/m^2^/s (low light LL; Fig. S14) before light intensity was increased for 45 min to 120 µmol/m^2^/s (high light HL: photoinhibition) and put back to 30 µmol/m^2^/s (LL: recovery). Values are expressed as the percentage of each culture’s yield before photoinhibition (T_0_). Means (± standard deviations) of triplicates from independent cultures are shown. Cell density for control (i.e. empty vector transgenic), Ot13-OE, Ot10-OE, Ot05-OE was in average, 48, 44, 32 and 48.10^6^ cell/ml respectively.

Thylakoid membrane PUFA are known to play a role in the regulation of photosynthetic processes (Allakhverdiev *et al*., 2009). We therefore investigated photosynthetic parameters of Δ6-DES-OE. Nevertheless, no significant changes regarding either photosynthesis efficiency or photoinhibition responses occurred under our conditions (Fig. 6C, S14)

## DISCUSSION

Regulation of C18-PUFA desaturation is required for the FA-profile remodelling of structural lipids in response to chilling in plants and cyanobacteria (Los *et al*., 2013); downstream synthesis of ω-3 and ω-6 VLC-PUFA in animals, fungi and microalgae also involves the fine-tuning of C18-PUFA amount notably by Δ6-DES. On the other hand, all front-end Δ6-DES studied so far are demonstrated, or assumed to be located in the ER (Meesapyodsuk & Qiu, 2012). By unveiling the occurrence of plastidic Δ6-DES with distinct substrate selectivity in the ancestral green picoalga *O. tauri* and of putative homologs in Chromista, our results strongly suggest the requirement of an autonomous control of plastidic C18-PUFA in several microalgae species. The entangled substrate features instructing the activity of these two novel Δ6-DES, the possible PUFA fluxes unveiled by Δ6-DES overexpression, as well as the physiological significance of pΔ6-DES are discussed.

### Substrate specificity of *O. tauri* Δ6-DES

DES specificity relies on intricate substrate features, including the acyl-chain position, length, unsaturation features, and the acyl-carrier nature (Heilmann *et al*., 2004a; Li, D *et al*., 2016). Domain swapping between DES of distinctive specificity, together with further amino acid mutation, could highlight the primary importance of His-Box and surrounding region for front-end desaturase activity and substrate specificity (Song *et al*., 2014; Li, D *et al*., 2016; Watanabe *et al*., 2016). However, neither the exact molecular features underlying DES (regio)specificity nor the hierarchical importance of the substrate features are yet clearly identified. Furthermore, assaying plant DES activity in *S. cerevisiae* might have introduced some bias by favoring the activity of microsomal desaturases and/or hampering the proper analysis of plastidic desaturases specificityin the absence of plastidic substrate (Heilmann *et al*., 2004b).

In this work, four different hosts were used to characterize pΔ6-DES substrate specificity. Lipid changes occurring in each of these organisms reflect a steady-state arising from both desaturation and overall FA fluxes. Nevertheless, comparison of lipidic features triggered by overexpression of each Δ6-DES in a given host and of the same Δ6-DES in different hosts allowed to gain insight into pΔ6-DES substrate specificity. In 16:3-plants, the Kennedy pathway contributes to the synthesis of plastidic lipid synthesis yielding 18:3/18:3-lipid species besides *sn-1/sn-2* 18:3/16:3n-3 (Browse *et al*., 1986b). In *O. tauri*, as in cyanobacteria and other most green microalgae, plastidic lipids correspond to *sn-1/sn-2* 18:X/16:X species. In contrast, *O. tauri* extraplastidic lipids encompass *sn-1/sn-2* SFA/16:4, VLC-PUFA/16:4 and di-homo-VLC-PUFA as major species (Degraeve-Guilbault *et al*., 2017). These distinctive positional signatures strongly suggest that, in *O. tauri*, plastidic lipids synthesis is independent of ER synthesis. On the other hand, acyl-lipid remodelling of plastidic lipids is assumed to be absent in *Synechocystis* PCC 6803, for which no acyl-turnover was ever reported (Ohlrogge & Browse, 1995). Concerning microalgae, MGDG has been proposed, yet not clearly proven, to be a platform of FA exchange supporting the incorporation of plastidic FA into TAG *in C. reinhardtii* (Li *et al*., 2012; Kim *et al*., 2018). The interpretation of our results takes support of this knowledge.

### Head-group specificity

As major changes occurred in galactolipids independently of the host, it can reasonably be concluded that at least MGDG is a substrate of both Ot05 and Ot10. Most interestingly, compared to Ot10 overexpression, Ot05 overexpression in *O. tauri, Synechocystis* PCC 6803 and *N. benthamiana* triggered a greater/exclusive accumulation of Δ6-DES products in SQDG whereas Ot10 overexpression in *Synechocystis* PCC 6803 and *N. benthamiana*, led to a greater accumulation of 18:3n-6 in PG. Ot10 overexpression further led to the production of 18:2-PG in *Synechocystis*. Altogether, these results show that Ot05 displayed a broad specificity for plastidic substrates while Ot10 appeared to be selective for PG, at least in heterologous hosts, as well as of some galactolipid species in the native host (see below). Interestingly, the plastidic ω-3-DES FAD8 was shown to display a preference for PG compared to the closely related FAD7 in *A. thaliana* (Roman *et al*., 2015).

Concerning the impact of pΔ6-DES overexpression on structural extraplastidic lipids, several evidences support that it most likely arises from the export of overproduced PUFA rather than from the access of pΔ6-DES to extraplastidic substrates. Indeed, the absence of desaturation activity in yeast supports that PC is not an accurate substrate (Domergue *et al*., 2003). Moreover, the fact that *Ostreococcus* Δ5-DES and Δ4-DES, which natural substrates are DGTA and possibly PDPT, displayed a low activity in *S. cerevisiae* supports the importance of the head-group for front-end DES activity (Hoffmann *et al*., 2008; Ahmann *et al*., 2011). We therefore propose that the first level of substrate recognition of *O. tauri* pΔ6-DES relies on plastidic lipid head-group. Recently, co-crystallization of the Stearoyl-CoA DES with its substrate revealed that interaction between the DES and the acyl-carrier was indeed fundamental to orient the acyl-chain in the catalytic tunnel of the enzyme (Wang *et al*., 2015).

Acyl-CoA-Δ6-DES overexpression in *O. tauri* importantly altered FA-profile of TAG and that of structural lipids only moderately. In contrast, acyl-CoA-Δ6-DES overexpression in *N. benthamiana* resulted in the accumulation of Δ6-desaturation products in all phospholipids.These results are coherent with the acyl-CoA specificity of Ot13: in *O. tauri* the incorporation of unwanted acyl-CoA Δ6-DES products into TAG possibly circumvents the alteration of extraplastidic lipids whereas in *N. benthamiana* leaves acyl-CoA are likely preferentially incorporated into PC, and in part transferred to MGDG (see below).

### ω-3/ω-6 and 16:4-galactolipid species selectivity

The preference of Ot05 for ω-3-substrates was reflected by the important accumulation of 18:4n-3 at the expense of 18:3n-3 observed in Ot05-OE from all of the host organisms. Conversely, Ot10 appeared globally more selective for ω-6 substrates. Nevertheless, Ot10 seemed to further display a ω-3 specificity for 16:4-galactolipid in which a rise of 18:4n-3 was paralleled by a drop of 18:3n-3. The ω-3-desaturation of the overproduced 18:3n-6 in Ot10-OE could possibly be involved in the rise of 18:4n-3. However, one would expect a higher accumulation of 18:3n-6 in Ot10-OE compared to Ot05-OE if only this route was used, which is not the case. Indeed, the endogenous ω-3-desaturase appeared to compete efficiently with the Δ6–desaturation of 18:2n-6 to 18:3n-6 maintaining a high amount of 18:3n-3 in pΔ6-DES-OE; therefore, the specific 18:4n-3 accumulation in 16:4-galactolipids most likely involves the Δ6-desaturation of 18:3n-3 precursors in the Ot10-OE *O. tauri* line.

In summary our results suggest that in addition to a distinctive preference of Ot05 and Ot10 for ω-3 substrates, Ot10 is further able to accept highly unsaturated galactolipids ω-3 substrates. Compared to *Synechocystis*, the higher activity of Ot10 on ω-6-substrates in *N. benthamiana* and *O. tauri* might also rely on the higher unsaturation degree of the non-substrate acyl-chain in these hosts. Therefore, the overall unsaturation degree of substrate molecular species appears to influence the activity/ ω-specificity of Ot10 while Ot05 activity seems independent of the unsaturation degree of the associated C16.

### Export of plastidic PUFA

Overexpression of each of the three Δ6-DES in *O. tauri* led to a similar increase of 20:4n-6 in DGTA and TAG and a specific increase of 18:4n-3 in DGTA for Ot05-OE. We previously reported that the acyl-CoA pool was enriched in 18:3n-3 and 18:4n-3, whose amounts were varying according to the plastidic C18-PUFA content (Degraeve-Guilbault *et al*., 2017). Together with the discovery of pΔ6-DES, these previous observations support that Δ6-desaturation products are exported from the plastid to the acyl-CoA pool. Though it cannot be excluded that pΔ6-DES have (limited) access to extraplastidic substrates as suggested for the plastidic ω-3-DES *C. reinhardtii*, front-end DES specificity is known to be more restricted than those of ω3-DES (Meesapyodsuk & Qiu, 2012; Nguyen *et al*., 2013; Wang *et al*., 2013). Unexpectedly, pΔ6-DES-OE impacted extraplastidic lipids to a much greater extent in *N. benthamiana*. As mentioned above, these changes are likely arising from reallocation of xenobiotic plastidic Δ6-desaturation products to other membranes, possibly to circumvent deleterious effects, and from the low capacity of leaves to synthesize TAG. In *A. thaliana* mutants deficient for the plastidic ω-3-DES (*FAD7*), alteration of 18:3 content in extraplastidic lipids has been reported and implied a two-way exchange of lipids between the chloroplast and the extrachloroplastic compartments (Browse *et al*., 1986a). Most of the following work focused on the transfer of lipid and PUFA from the ER to the chloroplast, establishing the idea that the reverse transport was negligible (Miquel & Browse, 1992; Li, N *et al*., 2016). Nevertheless, the recent characterization of two *Arabidopsis* plastid lipases mutants highlighted that plastidic PUFA indeed contributed to PUFA remodelling of extraplastidic lipid (Wang *et al*., 2017; Higashi *et al*., 2018).

### Co-regulation of DGDG and DGTA PUFA content

In *O. tauri* 16:4 is exclusively at *sn-2* position in both extraplastidic and plastidic lipids while being absent from the acyl-CoA pool and mostly at lateral positions in TAG. The SFA/16:4 combinations account for more than 70% and 20% of PDPT and DGTA species, respectively (Degraeve-Guilbault *et al*., 2017). In Ot05-OE, the concomitant 18:4n-3 increase and 16:4n-3 decrease specifically observed in DGDG and DGTA again rises the question about the origin of 16:4 extraplastidic again; indeed, the decreases of 20:4/16:4 DGTA and 18:3/16:4 DGDG could be related assuming that 16:4-DGDG species would yield DAG precursors for 16:4-extraplastidic species synthesis. Alternatively, the TAG pool could be involved. For instance, the peculiar TAGs species 16:4/16:4/18:X possibly gives DAG precursors with a 16:4-*sn-2*. Galactosyl-Galactosyl-Galactolipid-transferase, yet not identified in microalgae, supports the production of DAG from DGDG in plants under freezing conditions (Moellering *et al*., 2010). Export of specific DGDG-species to extraplastidic membranes has been reported for plants and microalgae under phosphate starvation (Jouhet *et al*., 2004; Khozin-Goldberg & Cohen, 2006).

### Significance of plastidic Δ6-DES

The absence of alteration in growth or photosynthetic processes in *O. tauri* Δ6-DES-OE is possibly related to compensatory mechanisms, including the increase of 16:0 and the decrease of 16:4. Photosynthesis defects were not detected in *Synechocystis* mutant devoided of 18:3n-3 and 18:4n-4, and could be unveiled only in very specific conditions in the *Arabidopsis* mutant lacking trienoic PUFA (Gombos *et al*., 1992; Vijayan & Browse, 2002). There is overall only little evidence that plastidic PUFA support photosynthetic processes (Mironov *et al*., 2012; Kugler *et al*., 2019).

The only plastidial front-end desaturase so far described was a Δ4-DES from *C. reinhardtii* and its Cyt-b5 domain was shown to be active *in vitro* (Zauner *et al*., 2012). Our data further show that a functional Cyt-b5 is absolutely required for *O. tauri* pΔ6-DES activity in both *N. benthamiana* and *Synechocystis*. These results illustrate the very tight co-evolution of Cyt-b5 and desaturase domains (Napier *et al*., 2003). It further indicates that a redox partner different from the eukaryotic Cyt-b5 oxidoreductase is involved (Napier *et al*., 2003; Kumar *et al*., 2012; Meesapyodsuk & Qiu, 2012). One possible candidate is the Ferredoxin NADP^+^-reductase (Yang *et al*., 2015).

The requirement of three Δ6-DES in an photosynthetic organism displaying the most reduced set of genes, points to the necessity of regulating the chloroplast and the cytosolic C18-PUFA pools distinctively. The existence of putative homologs of pΔ6-DES in other microalgae species, such as haptophytes and dinoflagellates, might also be related to the co-occurrence of 18:5 in galactolipids and the prevalence of VLC-PUFA in extraplastidic lipids. For the diatom *P. tricornutum*, the putative pΔ6-DES homolog might be involved in the desaturation of plastidic C16-PUFA as it had been suspected long ago (Domergue *et al*., 2002). For most *Mamiellophyceae*, with the exception of *Bathycoccus prasinos*, the two pΔ6-DES most likely arise from gene duplication. Ot10 would have evolved to restrict its specificity to a particular set of substrates. Transcriptional regulation of Ot05 by phosphate deprivation indicates that it is a physiological target under these conditions. The selectivity of Ot05 for SQDG, which is considered as a surrogate of PG under phosphate limitation, supports the importance of Ot05 in these conditions. Ot10 and the putative ω-3-DES transcriptional regulation might be targeted by other environmental cues, which still need to be discovered.

## METHODS

All chemicals were purchased from Sigma Chemical (St. Louis, MO, USA), when not stated otherwise.

### Biological material & cultures

*O. tauri* (clonal isolate from OtH95) was grown as previously described in artificial sea-water with either 5 µM or 35 µM NaH_2_PO_4_; penicillin (0,5 mg/ml) and streptomycin (0.25 mg/ml) together with centrifugation cycles (1000g, 5min) were used to reduce bacterial contamination; flow cytometry was used to assess growth and bacterial contamination (Degraeve-Guilbault *et al*., 2017). *Synechocystis* PCC 6803 was grown accordingly to (Kotajima *et al*., 2014). *N. benthamiana* plants were cultivated in a greenhouse under controlled conditions (16h:8h photoperiod, 25°C). *Agrobacterium tumefaciens* strain GV3101 was grown in Luria Broth medium at 30°C; *Saccharomyces cerevisiae* strain InvSc1 (*MATa adc2-1 his3-11,15 leu2-3,112 trp1-1 ura3-52*; Invitrogen) was grown in synthetic dextrose medium (5 mL, 50-mL Erlenmeyer flask, 30 °C, 180 rpm).

### Cloning strategy

PCR amplifications of DES ORF were achieved using Q5® Polymerase by two-step PCR on cDNA matrix (primers Table S4). Monarch DNA Gel Extraction kit was used when necessary (New Englands Biolabs, Iswitch, MA, US). Overexpression vectors were the pOtoxLuc for *O. tauri* (Moulager *et al*., 2010), pTHT2031S vector for *Synechocystis* PCC 6803 (Kotajima *et al*., 2014), the GATEWAY destination vectors pVT102-U-GW for *S. cerevisiae* (Domergue *et al*., 2010) and PK7W2G2D for *N. benthamiana* (Karimi *et al*., 2002). For subcellular localization, the final destination vector was pK7YWG2 (Karimi et al., 2002) N-terminal-YFP-fusion was used. Subcloning was achieved in pGEMT vector by restriction enzymes (Promega, Madison, WI, US), in pUC57 for codon-optimized sequence used in *Synechocystis* and for *S. cerevisiae* (GenScript Biotech, Netherlands) and/or in pDONR 221 for GATEWAY cloning. Restriction cloning was used for cloning in pOtLux, and ligation was used to introduce the synthetic gene in pTHT2031S In-Fusion® HD cloning kit (Takara Bio, Kusatsu, Japan). Sequencing was achieved by Genwiz (Genwiz, Leipzig, Germany).

Site-directed mutagenesis H >A of the HPGG motif of the Cyt-b5 domain was either performed by Genescript *(N. benthamiana*) or using In-Fusion® HD cloning kit (Takara Bio, Kusatsu, Japan) for *Synechocystis* after amplification using two mutagenic complementary primers for amplifying pTHT2031-*Ot5H46A*-S and pTHT2031-*Ot10H20A*-S from pTHT2031-Ot5-S and pTHT2031-Ot10-S, respectively. The mutated DNA sequence was validated for the correct modification using BigDye® Terminator v.3.1 (Life Technologies, Carlsbad, CA, US).

### RNA and cDNA preparation and quantitative RT-PCR analysis

RNeasy-Plus Mini kit (Qiagen, Hilden, Germany) was used for RNA purification; DNase I was used to remove contaminating DNA (DNA-free kit, Invitrogen, Carlsbad, USA) and cDNA obtained using the reverse transcription iScript™ supermix kit (Bio-Rad, Hercules, CA, USA). Real-time RT quantitative PCR reactions were performed in a CFX96™ Real-Time System (Bio-Rad) using the GoTaq^®^ qPCR Master mix (Promega, Madison, WI, USA) (Primers Table S4). Bio-Rad CFX Manager software was used for data acquisition and analysis (version 3.1, Bio-Rad). Ct method was used to normalized transcript abundance with the references mRNA *EF1α* (elongation factor), *CAL* (calmodulin), and *ACTprot2* (Actin protein-related 2). PCR efficiency ranged from 95 to 105%. Technical triplicate was used, and at least two independent experiments were achieved.

### Genetic transformation

*O. tauri* electroporation was adapted from (Corellou *et al*., 2009). Transgenics were obtained by electroporation and pre-screened accordingly to their luminescent level (Moulager *et al*., 2010). *S. cerevisiae* was transformed using a PEG/lithium acetate protocol, and FA supplementation was achieved as previously described (Dohmen et al, 1991). Control lines are transgenics of empty vectors. *N. benthamiana* leaves from five-week old plants were infiltrated with *Agrobacterium tumefaciens* previously transformed by electroporation; the p19 protein to minimize plant post-transcriptional gene silencing (PTGS) was used in all experiments (Voinnet *et al*., 2003). Briefly, *A. tumefaciens* transformants were selected with antibiotics (gentamycin 25µg/mL with spectinomycin 100µg/mL or kanamycin 50µg/mL). *Agrobacterium* transformants were grown overnight, diluted to a OD600 to 0.1 and grown to a OD600 of 0.6-0.8. Cells were re-suspended in 5 mL sterilized H_2_O for a final OD of 0.4 and 0.2 for overexpression and subcellular localization experiments, respectively and 1 mL was agroinfiltrated. Plants were analyzed 2 and 5 days after *Agrobacterium* infiltration for subcellular localization experiments and for overexpression, respectively.

*Synechocystis* transformation was achieved by homologous recombination (Williams, 1988). Briefly, the plasmid was transformed into ten-time concentrated cells of the Δ*desD* strain collected at mid-log phase. Subsequently, the cell was incubated at 30 °C under white fluorescent lamps for 16-18 hr and selected by 25 µg/mL chloramphenicol and 5 µg/mL spectinomycin on BG-11 solid media (1.5% w/v Bacto-agar).

### Lipid analysis

For all organisms, FA analyses and for *O. tauri* further lipid analysis were achieved accordingly to (Degraeve-Guilbault *et al*., 2017). Organic solvents all contained butylhydroytoluene as an antioxidant (0.001%). For *N. benthamiana* frozen material (one leaf broken into pieces) was preincubated in hot isopropanol (3ml, 75°C, 15 min, for PLD inhibition), further extracted with CHCl_3_ (1mL, Ultra-turax T25); Phase separation occurred upon addition of NaCl 2.5% (2000g, 10 min). Pellet was re-extracted (3 mL CHCl_3_: CH_3_OH 2:1 v/v); Organic phases were washed twice with 0.25v of CH_3_OH:H_2_O (10:9 v/v). The lipid extract was evaporated under a nitrogen stream and resuspended in CHCl_3_:CH_3_OH 2:1, v/v (200 µL). Lipids were separated by HP-TLC and chloroform/methanol/ glacial acetic acid/water (85:12:12:1 v/v/v/v). For *Synechocystis* (30mg DW, 3 mL CHCl_3_:CH_3_OH 2:1 v/v) extraction was achieved using glass bead vortexing. HP-TLC developments were achieved in the ADC2-chamber system (CAMAG) accordingly to (Degraeve-Guilbault *et al*., 2017) except for *Synechocystis* polar lipids (CHCl_3_:CH_3_OH/CH_3_COOH/H2O 85:12:12:1 v/v/v/v) (Sallal *et al*., 1990). Lipids were visualized and collected as previously described.

MS analyses of *O. tauri* glycerolipid species lipids was performed accordingly to the method describe previously (Abida *et al*., 2015). Purified lipids were introduced by direct infusion (electrospray ionization-MS) into a trap-type mass spectrometer (LTQ-XL; Thermo Scientific) and identified by comparison with standards. Lipids were identified by MS^2^ analysis with their precursor ion or by neutral loss analyses. Positional analysis of FA in glycerolipids was achieved as previously described (Degraeve-Guilbault *et al*., 2017).

### Confocal microscopy

Live cell imaging was performed using a Leica SP5 confocal laser scanning microscopy system (Leica, Wetzlar, Germany) equipped with Argon, DPSS, He-Ne lasers, hybrid detectors, and 63x oil-immersion objective. *N. benthamiana* leave samples were transferred between a glass slide and coverslip in a drop of water. Fluorescence was collected using excitation /emission wavelengths of 488/490-540 nm for chlorophyll, 488/575-610 nm for YFP, and 561/710-740 nm for m-cherry. Colocalization images were taken using sequential scanning between frames. Experiments were performed using strictly identical confocal acquisition parameters (*e*.*g*. laser power, gain, zoom factor, resolution, and emission wavelengths reception), with detector settings optimized for low background and no pixel saturation.

### Photosynthesis measurement

Measurements were made using a PhytoPAM (Heinz Walz GmbH, Germany).

### Light-response of photosystem II activity

Rapid light-response curves (RLCs) of chlorophyll fluorescence of the cultures were achieved accordingly (Serodio *et al*., 2006). Briefly, the cultures were exposed to 12 increasing actinic light levels (10-s light steps of 100 µE increase from 64 to 2064 µE), and the electron transport rates (ETR) were calculated on each step to draw RLCs. The following parameters were extracted from the ETR-irradiance curve fitted to the experimental data: the initial slope of the curve (*α*), the light-saturation parameters (I_k_), and the maximum relative electron transport rate (ETRmax).

### Photoinhibition experiment

Optimal conditions for photosystem II inhibition and recovery were adapted from (Campbell & Tyystjarvi, 2012). Cultures (50 mL, triplicate) were maintained under fluorescent white light (low light: 30.4±1.0 µE, white light) without agitation at 20.2±0.2 °C and moved to high light (117.6±4.9 µE, blue LED) for 45 min (photoinhibition); photorecovery under initial condition was monitored for over 2 h. One mL sampling was used to assess photosynthetic efficiency (quantum yield of photochemical energy conversion in PSII; Y corresponds to Yield = dF/Fm).

### Sequences analyses

DES domain-containing sequences were retrieved from genomic and transcriptomic data from NCBI (Bioproject Accession: PRJNA304086 ID: 304086). Annotated ORF were manually checked for the completion of Nt sequences in species from the class Mamiellalophyceae; cTP were predicted from PredAlgo (Tardif et al., 2012); alignment of Mamiellalophyceae homologous sequences was used to further determine putative cTP assumed to correspond to the non-conserved Nt region (Snapgene trial version, Clustal omega). These non-conserved regions were discarded for expression in *S. cerevisiae* and *Synechocystis*. Codon-optimized sequences were obtained from Genewiz (Europe).

## Accession numbers

[At]S8: Arabidopsis thaliana AEE80226.1, [Bo]6: Borago officinalis AKO69639.1, [Ce] Caenorabditis elegans CAA94233.2, [Cr]4p Chlamydomonas reinhardtii AFJ74144.1, [Ep]6 Echium plantagineum AAZ08559.1, [Eg]8 Euglena gracilis AA D45877.1, [Eh] Emiliana huxleyi putative protein XP_005793257.1, [Gt] Guillardia theta CCMP2712 putative protein XP_005823787.1, [Ig]5/6 Isochrisis galbana ALE15224.1 and AHJ25674.1, [Lb] Leishmania brazilensis putative protein XP_001569342.1, [Li]6 Lobosphaera incisa ADB81955.1, [Ma]6 Mortierella alpine BAA85588.1, [Mp]6CoA Micromonas pusilla XP_003056992.1, [Ms]6CoA Mantoniella squamata CAQ30479.1, [No]6 Nannochloropsis oculata, ADM86708.1, [Ot] Ostreococcus tauri CAL56435.1 (Δ6 acyl-CoA-DES Ot13.1), CEF97803.1 (Ot05), CEF99426.1 (Ot10); CEG01739.1 (Ot15); CEF99964.1, (D5-DES), CEF96519.1 (4ER), CEG00114.1, (D4p, Ot13.2), [Pt] Phaeodactylum tricornutum XP_002185374.1 (putative protein), EEC45637.1 (D6-DES) EEC45594.1 (D5-DES), [Vc] Volvox carteri XP_002953943.1, [SS] Synechocystis sp BAA18502.1, [Tp] Thalassiosira pseudonana, XP_002289468.1 (putative protein) AAX14504.1 (S8); AAX14502.1 (8); AAX14505.1 (6), XP_002297444.1 (4).

## Acknowledgements

This work was supported by the Région-Aquitaine grant “omega-3” and the University of Bordeaux grant Synthetic biology SB2 “Pico-FADO”. Routine lipids analyses were performed at the Metabolome Facility of Bordeaux-MetaboHUB (ANR-11-INBS-0010). Imaging was performed at the Bordeaux Imaging Center, member of the national infrastructure France BioImaging.

